# Strong effects of parasite genotype on drug susceptibility in the Indian subcontinent

**DOI:** 10.1101/2022.08.17.504263

**Authors:** Cooper Alastair Grace, João Luís Reis-Cunha, Daniel C. Jeffares

## Abstract

Intracellular parasites of the *Leishmania donovani* species complex cause visceral leishmaniasis (VL). For parasitic diseases, VL has a mortality rate second only to malaria, and is associated with poverty-stricken areas of the world: primarily Brazil, East Africa and the Indian subcontinent (ISC). Miltefosine (MIL) and the antimonal sodium stibogluconate (SSG) are drugs used in the treatment of leishmaniasis. However, treatment efficacy is variable, and the numbers of reports of parasite resistance to both drugs have risen since their introductions, particularly in the ISC. To assess the level of parasite genotype contribution to drug resistance, we utilised the sequencing and associated drug susceptibility data from Imamura *et al*. (2016) to estimate heritability and GWAS using LDAK. We obtained strong heritability results, with mainly SNP/indel variations associated with SSG and copy number variants associated with MIL resistance, respectively. However, GWAS results were inconclusive, suggesting that, although the parasite genotype directly influences drug resistance, the effect might be multifactorial.

## Introduction

*Leishmania* are intracellular parasitic protozoans, spread by sandfly vectors, which cause leishmaniasis, a collection of neglected tropical diseases. Species of the *Leishmania donovani* species complex (LDSC), which include *L. donovani* and *L. infantum*, are the main causative agents of visceral leishmaniasis (VL), the second largest parasitic cause of death after malaria (Alvar *et al*., 2012). The major endemic populations of East Africa, the Indian subcontinent (ISC) and Brazil are genetically divergent from one another and elicit diverse clinical outcomes (Imamura *et al*., 2016; Alves *et al*., 2018; Siriwardana *et al*., 2019; Franssen *et al*., 2020; Grace *et al*., 2021).

Miltefosine (MIL), currently the only oral treatment for VL, was introduced in India around 20 years ago, with high treatment success rates (Jha *et al*., 1999; Sundar *et al*., 2002); however drug resistance arose within a decade (Rijal *et al*., 2007, 2013; Sundar *et al*., 2012). The mechanism for resistance is unclear outside of Brazil, where a deletion of ∼12kb on chr31, containing four genes, is strongly associated with MIL resistance (Carnielli *et al*., 2018). Alternative mechanisms for MIL resistance have been proposed (Pérez-Victoria *et al*., 2003; Rakotomanga, Saint-Pierre-Chazalet and Loiseau, 2005; Cojean *et al*., 2012; Hefnawy *et al*., 2017; Van Bockstal *et al*., 2020).

An alternative treatment for VL, sodium stibogluconate (SSG) is a pentavalent antimonial with an ever increasing rate of failure, particularly in the ISC (Sundar *et al*., 2000; Rijal *et al*., 2003), attributed to parasite SSG resistance (Croft, Sundar and Fairlamb, 2006; Rijal *et al*., 2007; Downing *et al*., 2011; Vanaerschot *et al*., 2012). Several mechanisms for SSG resistance have been uncovered (Hefnawy *et al*., 2017), including copy number changes in the gene MRPA, responsible for sequestration of thiol complexes (Mittal *et al*., 2007; Imamura *et al*., 2016).

Imamura *et al*. (2016) evaluated potential genomic variations associated with SSG resistance in *L. donovani* isolates from the ISC. The authors found no statistically significant associations between any specific variants and SSG or MIL resistance. They proposed a potentially multifactorial origin for SSG resistance, including a frameshift insertion in the aquaglyceroporin-1 gene and higher copy numbers of the H-locus and MRPA gene. However, they did not assess the heritability - the proportion of phenotype that is attributable to genotype - of drug susceptibility in the analysed isolates.

Here, we apply genome-scale quantitative genetic analyses to investigate the effects of parasite genotype on MIL and SSG susceptibility present in ISC isolates of *L. donovani* sequenced by Imamura *et al*. (2016). Our analyses suggest that susceptibility to both drugs is strongly heritable, with SSG resistance being mainly attributed to SNPs and MIL to gene copy number variants. Due to low nucleotide diversity, low number of samples and a potentially multifactorial origin of these traits, specific genetic variants associated with susceptibility cannot be recovered from these data.

## Methods

We utilised the genome sequencing and associated drug susceptibility data produced by Imamura *et al*. (2016) of ISC isolates (*n*=229). Sequencing reads in Bioproject ERP000140 were downloaded from ENA and mapped onto the *L. donovani* BPK282A1 v.46 (https://tritrypdb.org) reference genome using bwa v.0.7.17 (Li and Durbin, 2009) with default parameters. Read duplicates were removed with SAMtools v.1.9 (Li *et al*., 2009). Copy number variants (CNVs) were determined using resulting BAM files, which were first filtered to retain mapping quality of ≥30. Using BEDtools (Quinlan, 2014) *genomecov*, the mean coverage of each gene (*n*=8,005) was estimated, and normalised by total genome coverage.

For each strain, SNPs and indels were called using Genome Analysis Toolkit (GATK) *HaplotypeCaller* v.4.1.0.0 (Depristo *et al*., 2011) and the ‘discovery’ genotyping mode of Freebayes v.1.3.2 (https://github.com/ekg/freebayes), only accepting calls discovered by both methods. Isolate VCFs were merged into a single population VCF with BCFtools v1.9 (Danecek *et al*., 2021). We retained only biallelic variants, with read depth within 0.3-1.7x the chromosome coverage, excluding any variants called on repetitive regions, as described in Grace *et al*. (2021). Further filtering removed sites with any of the following: DPRA < 0.73 or > 1.48; QA or QR < 100; SRP or SAP > 2000; RPP or RPPR > 3484; PAIRED or PAIRPAIREDR < 0.8; MQM or MQMR < 40. As chromosome 31 is generally supernumerary, we specified DPB < 30401 or > 121603 to be removed, and for remaining chromosomes, DPB < 18299 or > 73197 (<0.5x or >2x median DPB). Biallelic indels were filtered to remove sites with any of the following: DPRA < 0.73 or > 1.48; QA or QR < 100; SRP or SAP > 2000.

Linkage disequilibrium (LD) decay was assessed with the package PopLDdecay (Zhang *et* al., 2019). However, due to low nucleotide diversity, low recombination rates and high isolate similarity, a typical decay pattern was not observed, as r^2^ ≅ 1.0 for >300kb. SNPs were evenly sampled before plotting with R (R Core Team, 2022).

We considered MIL and SSG susceptibility as phenotypic ‘traits’ and used SNPs/indels and CNVs as parasite genotypes to estimate the total effect of parasite genetic variation on the traits (narrow sense heritability), using LDAK v.5.1 (Speed, Holmes and Balding, 2020). From the 229 isolates, 50 were assessed for SSG susceptibility, and 92 for MIL susceptibility (Imamura *et al*., 2016). Drug susceptibility data were normalised (0-1) prior to analysis.

PLINK v1.9 (Chang *et al*., 2015) was used to create the required binary files from the VCF. The heritability contribution of genotype class (SNPs/indels and CNVs) to each trait was estimated using LDAK restricted maximum likelihood (REML). GWAS was used to identify variants that were statistically associated with traits. The LDAK option --linear was used to conduct mixed model GWAS, using the kinship matrix derived from SNP/indel variants to control for unequal strain relatedness. Significance thresholds were determined via trait permutation. For both traits, values were permuted 1000 times, and parasite genotype-drug susceptibility trait associations estimated. The lowest *P* value was determined from each replicate permutation analysis, and the 1st and 5th percentiles used as the significance threshold for associations (i.e. 1% and 5% family-wise false discovery rate) for each trait.

## Results and Discussion

We identified 35,382 SNPs and indels across the 229 isolates used, based upon the *L. donovani* BPK282A1 reference genome. Gene copy number variants (CNVs) values ranged from 0.357-5.676.

We determined narrow sense heritability for MIL and SSG susceptibility for both SNPs/indels and CNVs (**Figure 1**). SSG susceptibility is a complex trait (total heritability 92%, SD ±7%), with contributions from both SNPs/indels (88%, SD ±11%) and minimally CNVs (4%, SD ±4%). MIL susceptibility is independently affected by CNVs (67%, SD ±16%), consistent with previous analyses (Carnielli *et al*., 2018). The deletion of the miltefosine sensitivity locus (MSL) which results in MIL resistance is however not observed outside the new world (Schwabl *et al*., 2021), indicating that this locus is not the reason for MIL susceptibility in isolates of the ISC.

**Figure 1.**
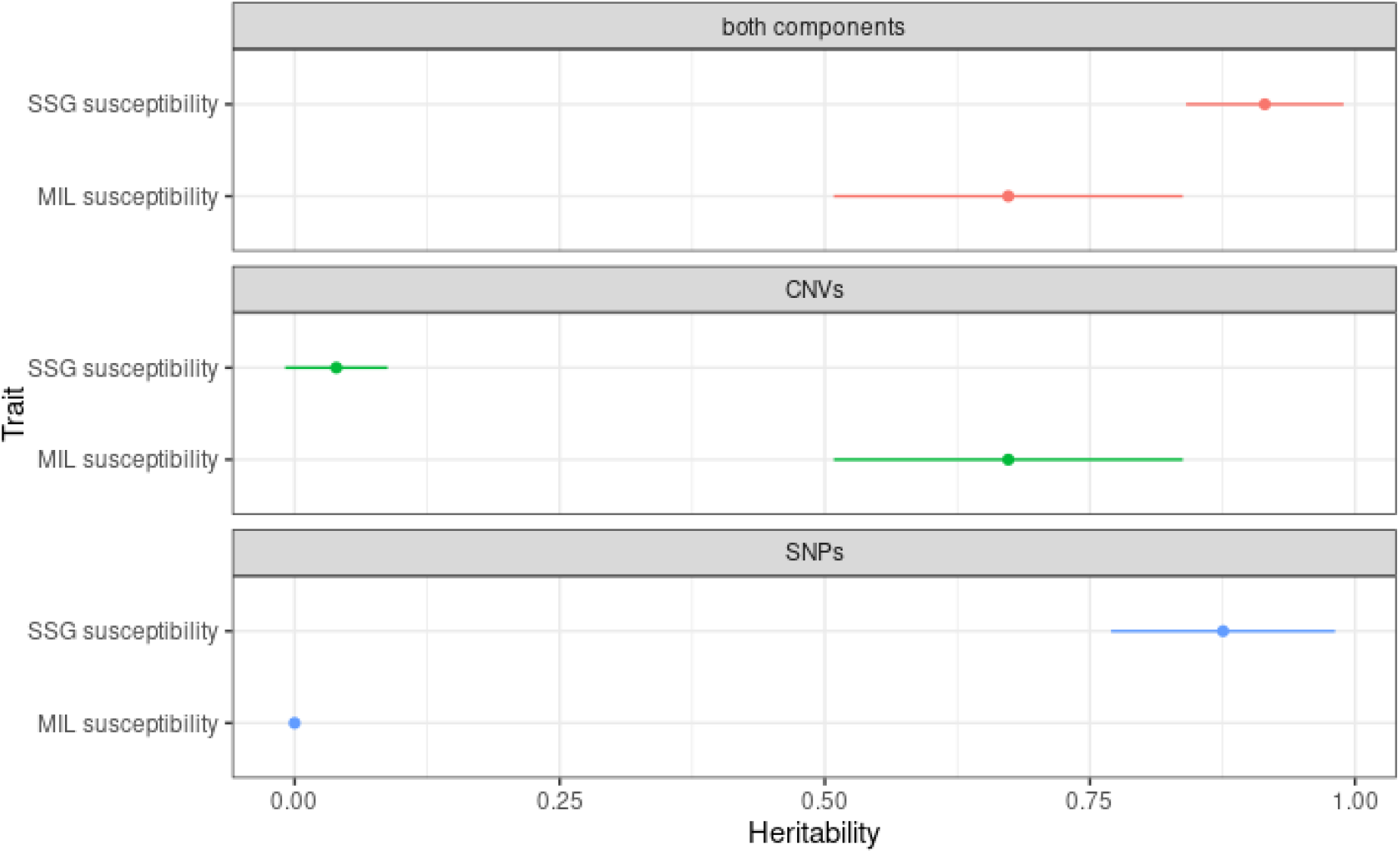
Heritability estimates using LDAK’s composite model. Analysis used susceptibility values which were normalised. Each point represents the estimated heritability value, with horizontal bars representing the standard deviation (+/-).

QQ-plots of *P* values for GWAS indicate there are multiple variants above the expected threshold (**Figure 2**). However, GWAS analysis did not recover significant variants associated with MIL or SSG susceptibility (**Figure 3**), indicating that multiple variants cumulatively contribute to parasite phenotype. We additionally assessed linkage disequilibrium (LD) decay within these isolates (**Figure 4**). Due to low nucleotide diversity (Grace *et al*., 2021), low recombination rates and highly similar isolates (Imamura *et al*., 2016), r^2^ values suggest that SNPs are strongly linked over large genetic distances. Together with population structure (Power, Parkhill and de Oliveira, 2017), these may be factors which have prevented the discovery of statistically significant variants associated with drug susceptibility.

**Figure 2.**
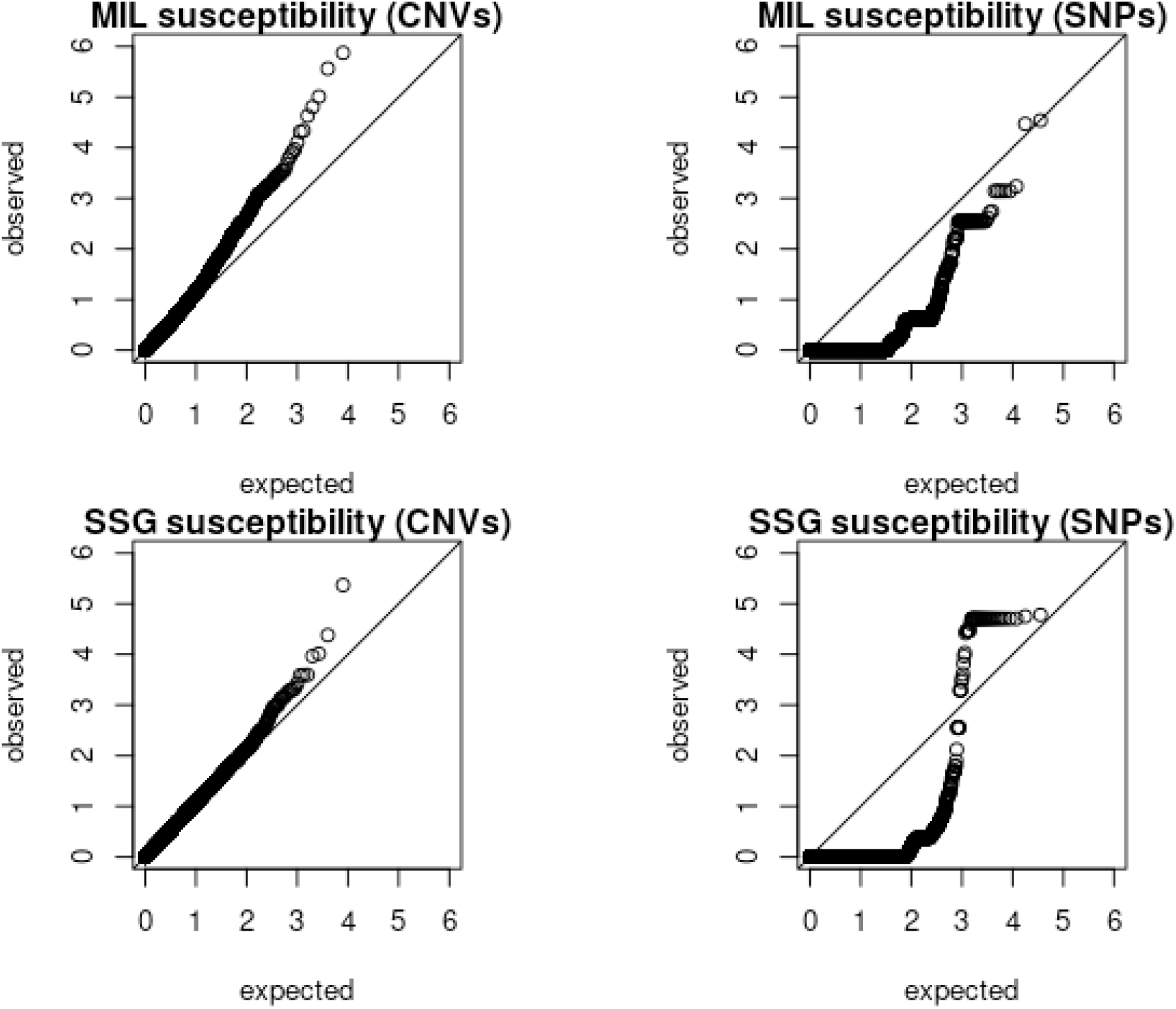
QQ-plots of observed and expected *P* values for GWAS analysis of SNPs and CNVs for MIL and SSG susceptibility. Top row: MIL susceptibility for CNVs and SNPs, respectively; bottom row: SSG susceptibility for CNVs and SNPs, respectively.

**Figure 3.**
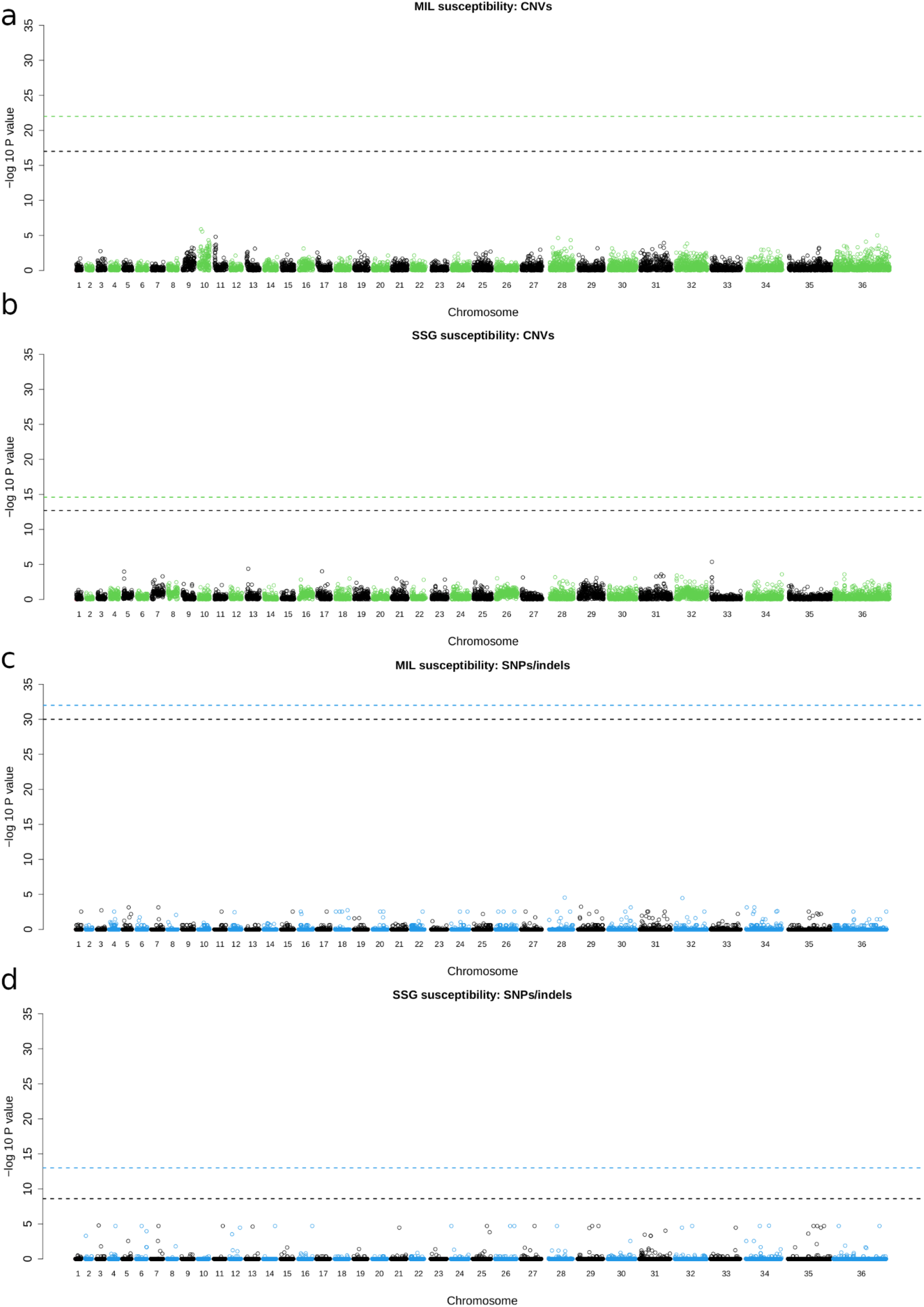
Manhattan plots for GWAS -log_10_ *P* values. No significant associations are recovered for either MIL or SSG susceptibility when using SNPs/indels or CNVs as genetic variants. Panels are: **a** - MIL susceptibility using CNVs; **b** - SSG susceptibility using CNVs; **c** - MIL susceptibility using SNPs/indels; **d** - SSG susceptibility using SNPs/indels. Horizontal lines indicate significance thresholds, determined by 1000 permutations, where green/blue represents the 99th quantile for CNVs and SNPs/indels, respectively, and black represents the 95th quantile. Note that the Y-axis scale is consistent across all panels.

**Figure 4.**
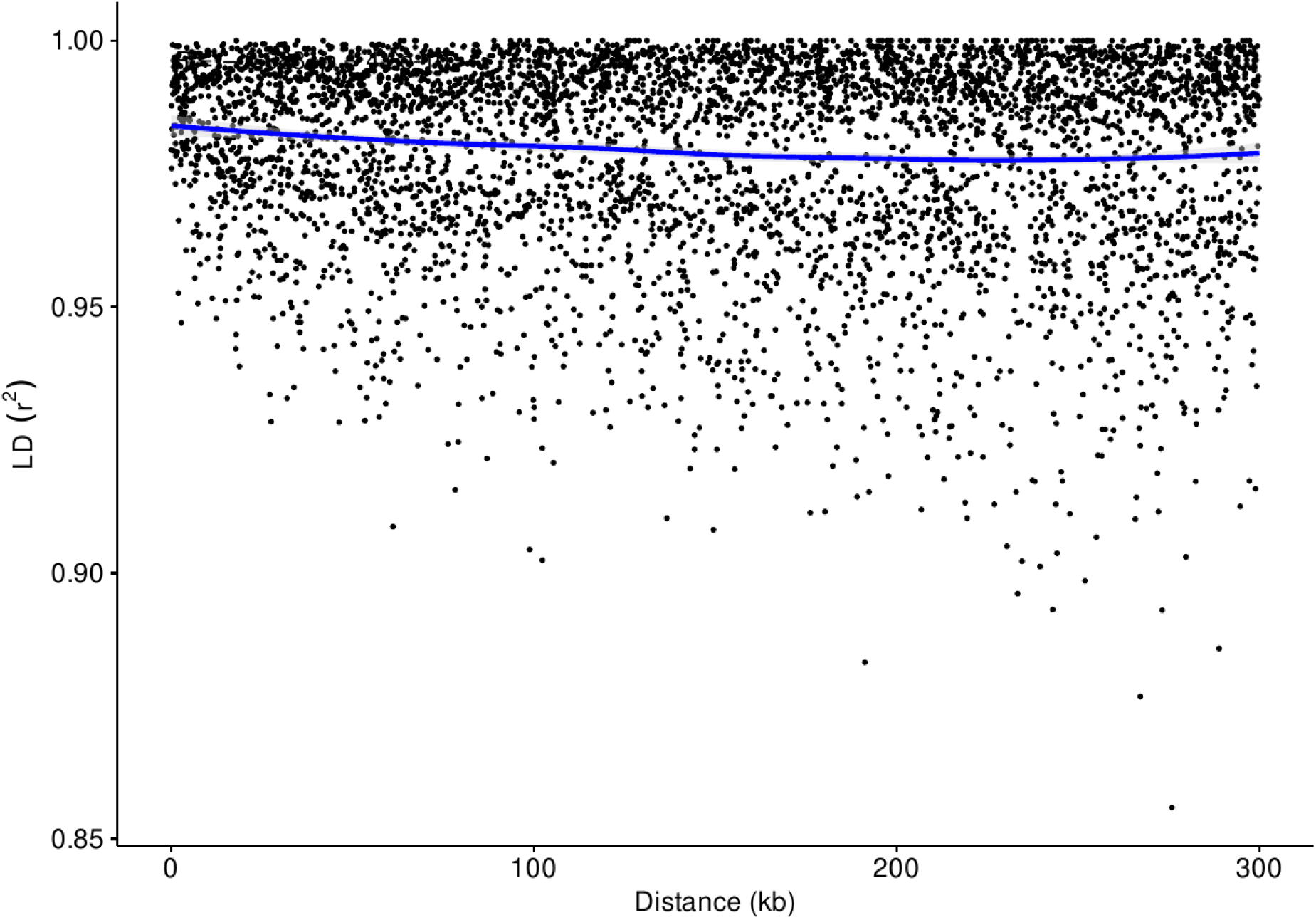
Linkage disequilibrium (LD) decay. Values were obtained with PopLDdecay (Zhang *et al*., 2019) using default parameters. SNPs were evenly sampled before plotting, with the full version available on FigShare. Low nucleotide diversity, low recombination rates and high similarity between isolates cause SNPs to appear linked over large genetic distances.

## Abbreviations

LDSC: *Leishmania donovani* species complex
ISC: Indian Subcontinent
MIL: miltefosine
SSG: sodium stibogluconate
VL: visceral leishmaniasis
LD: linkage disequilibrium
SNP: single nucleotide polymorphism
CNV: copy number variant
GWAS: genome-wide association study
REML: restricted maximum likelihood

## Data availability

All sequencing (BioProject ERP000140) and drug susceptibility data are publicly available. PDF versions of figures are available on FigShare: https://figshare.com/projects/Effects_of_parasite_genotype_on_drug_susceptibility_in_the_Indian_subcontinent/145485

## Author contributions

CAG - formal analyses, wrote and edited the manuscript.

JLRC - formal analyses, wrote and edited the manuscript.

DCJ - project conception, wrote and edited the manuscript.

